# Modulation of the canonical Wnt activity by androgen signaling in prostate epithelial basal stem cells

**DOI:** 10.1101/2020.01.10.902270

**Authors:** Yueli Liu, Jiawen Wang, Corrigan Horton, Sol Katzman, Tao Cai, Zhu A. Wang

## Abstract

Both the canonical Wnt signaling and androgen signaling are important factors regulating prostate organogenesis. How these two pathways crosstalk to regulate prostate stem cell functions remain unclear. Here, we show that while canonical Wnt activity is required for prostate basal stem cell multipotency in vivo, ectopic Wnt activity does not promote basal-to-luminal cell differentiation. We provide evidence that androgen signaling may keep Wnt activity in check. In prostate organoid culture from basal cells, dihydrotestosterone (DHT) antagonizes R-spondin-stimulated organoid growth in a concentration-dependent manner. Molecular analyses of organoids under different treatment conditions showed that androgen signaling down-regulated the expressions of a Wnt reporter as well as many Wnt target genes. Pathway analysis and gene set enrichment analysis of organoid RNA-seq data also revealed the canonical Wnt signaling as a key pathway distinguishing organoids treated with or without DHT. Notably, DHT treatment enhanced AR and β–catenin binding in the nuclei of prostate organoids, providing possible mechanistic clues. Our results reveal a critical role of AR signaling in modulating canonical Wnt activity in prostate basal cells to regulate their multipotency.

## Introduction

Signaling crosstalks are crucial for regulating adult stem cells in an organ. The canonical Wnt signaling pathway is a key player in stem cell self-renewal and differentiation in multiple tissues (Clevers et al., 2014; Holland et al., 2013). Its interaction with organ-specific hormone signals may be important for orchestrating the functions of adult stem cells in that tissue. In the prostate, both the canonical Wnt signaling and androgen signaling pathways have been shown to be essential in driving organogenesis (Toivanen and Shen, 2017). Specifically, stromal Wnt secretion promotes prostate bud branching morphogenesis and expression of the prostate-specific transcription factor Nkx3.1 in the epithelium (Francis et al., 2013; Julio et al., 2013; Simons et al., 2012). Classical tissue recombination experiments and conditional knockout experiments have revealed the role of stromal androgen receptor (AR) in instructing the specification of prostate epithelium through paracrine signals (Cunha et al., 1992; Lai et al., 2012). However, how Wnt and androgen signals interact to promote prostate organogenesis and whether their interaction affects prostate stem cells remain unknown.

Basal and luminal cells are the two major cell types lining the prostate epithelium. During prostate organogenesis, basal cells behave as stem cells to generate luminal cells and rare neuroendocrine cells (Ousset et al., 2012). Although basal stem cell multipotency become restricted in the mature prostate (Choi et al., 2012; Wang et al., 2013), isolated adult basal cells can be reactivated to generate prostatic tissues in the renal-grafting assay (Wang et al., 2014; Xin et al., 2003), and are more efficient in generating prostate organoids in cultures compared to luminal cells (Chua et al., 2014; Karthaus et al., 2014). The organoid culture medium contains R-spondin, an agonist of the canonical Wnt pathway (Binnerts et al., 2007; Carmon et al., 2011; de Lau et al., 2011), demonstrating a positive role of Wnt signaling for prostate organoid formation and expansion (Karthaus et al., 2014). On the other hand, conditional knockout of β-catenin in adult prostate basal cells appeared to negatively affect their luminal differentiation, as revealed by lineage tracing during a 6-week period (Lu and Chen, 2015). Recently, our lab performed long-term basal cell lineage tracing with AR conditional knockout, and demonstrated that AR within basal cells is required for basal-to-luminal cell differentiation in vivo (Xie et al., 2017). Despite these findings, how Wnt and androgen signaling pathways cooperate to promote basal cell multipotency is unclear. In this study, we show that ectopic Wnt activity leads to prostate basal cell over-proliferation, but not luminal differentiation, and that androgen signaling can down-regulate Wnt activity in basal cells, possibly through binding with β-catenin to interfere with target gene transcription.

## Results and Discussion

### Intermediate Wnt activity is essential for prostate basal-to-luminal cell differentiation

Although a previous lineage-tracing study suggested a role of canonical Wnt activity in promoting basal-to-luminal differentiation (Lu and Chen, 2015), the long-term effect of β-catenin deletion in basal cells is unknown. To this end, we utilized the *CK5-CreER*^*T2*^; *R26R-CAG-YFP/+* reporter for basal cell lineage tracing, for which we showed previously marks almost all prostate basal cells (Xie et al., 2017). We tamoxifen-induced *CK5-CreER*^*T2*^; *Ctnnb1*^*fl/fl*^; *R26R-CAG-YFP/+* adult male mice (termed *Bas*^*bcat−/−*^) and *CK5-CreER*^*T2*^; *R26R-CAG-YFP/+* controls (termed *Bas*^*WT*^), and performed long-term lineage tracing under homeostasis (Fig. 1A). Since β-catenin is a structural protein expressed in luminal cells contacting the basal layer, its loss in individual basal cells could not be easily distinguished by immunofluorescence (IF) staining (Fig. S1A). Nevertheless, efficient β-catenin deletion driven by *CK5-CreER*^*T2*^ was confirmed by flow-sorting of YFP+ prostate basal cells (Fig. S1B) for qRT-PCR analysis (Fig. S1C), and inferred by observing hair loss through time (Fig. S1D) (Kishimoto et al., 2000). Six months after induction, basal-to-luminal cell differentiation was significantly reduced in the *Bas*^*bcat−/−*^ group compared to *Bas*^*WT*^ (Fig. 1B, 1C), while basal cell proliferation rates were unchanged as measured by Ki67 staining and BrdU incorporation assay (Fig. 1D, 1E). These results are consistent with the previous 6-week tracing data in the Lu and Chen study. However, contrary to their finding in the tumor models (Lu and Chen, 2015), we found that β-catenin deletion did not rescue the progression of basal-derived *Pten*-null tumors, as *CK5-CreER*^*T2*^; *ctnnb1*^*fl/fl*^; *Pten*^*fl/fl*^; *R26R-CAG-YFP/+* tumors (*Bas*^*bcat−/−Pten−/−*^) were morphologically indistinguishable from those of *CK5-CreER*^*T2*^; *Pten*^*fl/fl*^; *R26R-CAG-YFP/+* (*Bas*^*Pten−/−*^) (Fig. 1F). The double knockout tumors still contain cribriform pattern and luminal differentiation (Fig. 1G), despite their lack of β-catenin expression (Fig. 1H). The cause of the discrepancy is unclear. We conclude that Pten loss can override the requirement of β-catenin in promoting basal-to-luminal cell differentiation.

**Figure 1.**
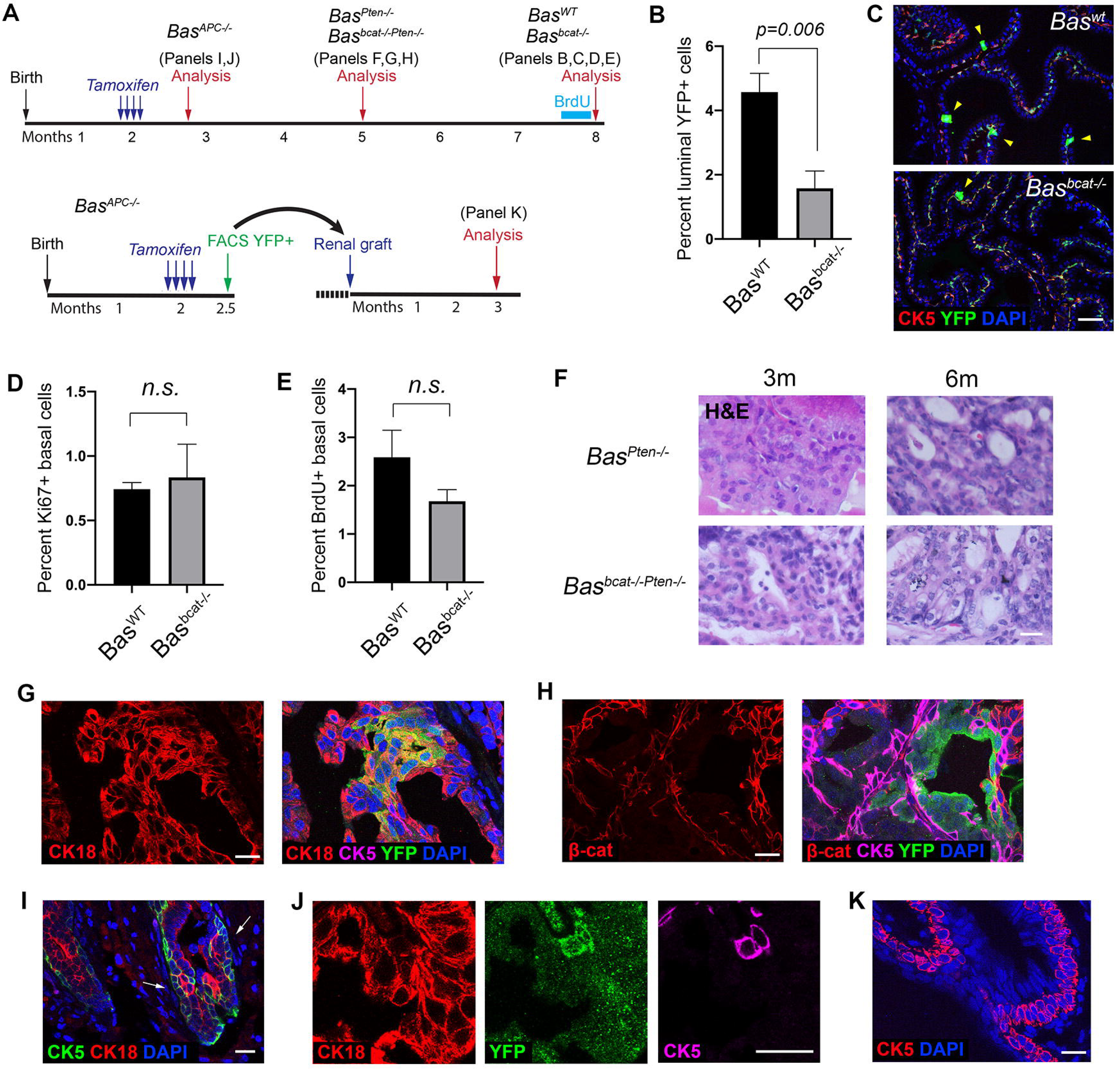
Lineage tracing analysis of basal cells of different levels of Wnt activity. (**A**) Timeline of experiments for lineage tracing and renal grafting of Wnt-mutant-related basal cells. (**B**) Percentage of YFP^+^ luminal cells decreased significantly in *Bas*^*bcat−/−*^ mice (N=5) compared to *Bas*^*WT*^ mice (N=4) at 6 months post induction by student’s t-test. (**C**) Representative IF images of YFP^+^ luminal cells (arrowheads) in the *Bas*^*WT*^ and *Bas*^*bcat−/−*^ prostate 6 months post tamoxifen induction. (**D, E**) Bar graphs showing proliferation rates in *Bas*^*WT*^ and *Bas*^*bcat−/−*^ basal cells by Ki67 (**D**) and BrdU (**E**). n.s., not significant by student’s t-test. (**F**) H&E staining of tissue sections from *Bas*^*Pten−/−*^ and *Bas*^*Pten−/−;Bcat−/−*^ mice at 3 months and 6 months post tamoxifen induction. (**G, H**) IF image showing increased basal-to-luminal differentiation (**G**) and loss of β-catenin (**H**) in *Bas*^*Pten−/−;Bcat−/−*^ tumor regions at 3 months post induction. (**I**) Squamous basal cell phenotype (arrows) seen in *Bas*^*APC−/−*^ mice at 2 weeks post induction. (**J**) IF staining showing the strict basal cell identity of the squamous-like cells. (**K**) IF showing *Bas*^*APC−/−*^ basal cell over-proliferation 3 months after renal grafting. Scale bars in **C** correspond to 50 μm, and in **F-K** to 20 μm. Error bars correspond to one s.d.

To assess whether gain of Wnt signaling activity could enhance basal-to-luminal cell differentiation, we tamoxifen-induced *CK5-CreER*^*T2*^; *APC*^*fl/fl*^; *R26R-CAG-YFP/+* mice (termed *Bas*^*APC−/−*^) and performed 3-week lineage tracing (Fig. 1A). We had to analyze the prostate morphology at this early time point due to mice succumbing to rapidly developing basal tumors growing elsewhere. While the majority of prostate epithelial cells appeared normal, we observed clones of basal cells stacked with each other, resembling squamous tumors (Fig. 1I). Those cells were strictly basal, and did not express the luminal marker CK18 (Fig. 1J). We then tried to propagate the basal-APC-deleted prostate tissues by renal grafting those cells onto immunodeficient mice (Fig. 1A), and again observed such basal cell over-proliferation phenotype after three months of growth (Fig. 1K). Taken together, these results suggest that Wnt signaling activity needs to be tightly controlled in vivo since intermediate Wnt activity is essential for basal-to-luminal cell differentiation.

### Ectopic Wnt activity enhances basal stem cell activities in organoid assay

We next investigated the effects of different Wnt activity levels on prostate basal stem cells using a defined organoid culture protocol (Drost et al., 2016). Wild-type and Wnt-activated basal cells were isolated from induced *Bas*^*WT*^ and *Bas*^*APC−/−*^ mice, respectively, and were cultured under three conditions: (1) “basement medium” (BM condition) containing all the factors as described in the protocol except for R-spondin and dihydrotestosterone (DHT), (2) BM plus R-spondin (R condition), and (3) BM plus the Wnt antagonist Dkk-1 (Dkk condition) (Glinka et al., 1998). Representative images of organoids growing for 7 days under different conditions are shown in Fig. 2A. These organoids typically exhibit CK5-expressing basal cells on the outskirt and CK18-expressing luminal cells in the center (Fig. 2A). Organoids of *Bas*^*APC−/−*^ cells were significantly larger than those of *Bas*^*WT*^ (Fig. 2B), and contained significantly more complex structures such as increased branching (Fig. 2C), suggesting ectopic Wnt activity enhances basal stem cell activities and organoid growth in vitro. Consistently, R-spondin treatment increased organoid size and branching morphogenesis compared to BM for both *Bas*^*APC−/−*^ and *Bas*^*WT*^ organoids, while Dkk-1 treatment had the opposite effects (Fig. 2B, 2C). Notably, robust organoid growth under the BM or R condition was achieved without adding DHT, a component described in the defined prostate organoid protocol (Drost et al., 2016), suggesting that DHT is not required for basal stem cell functions in vitro.

**Figure 2.**
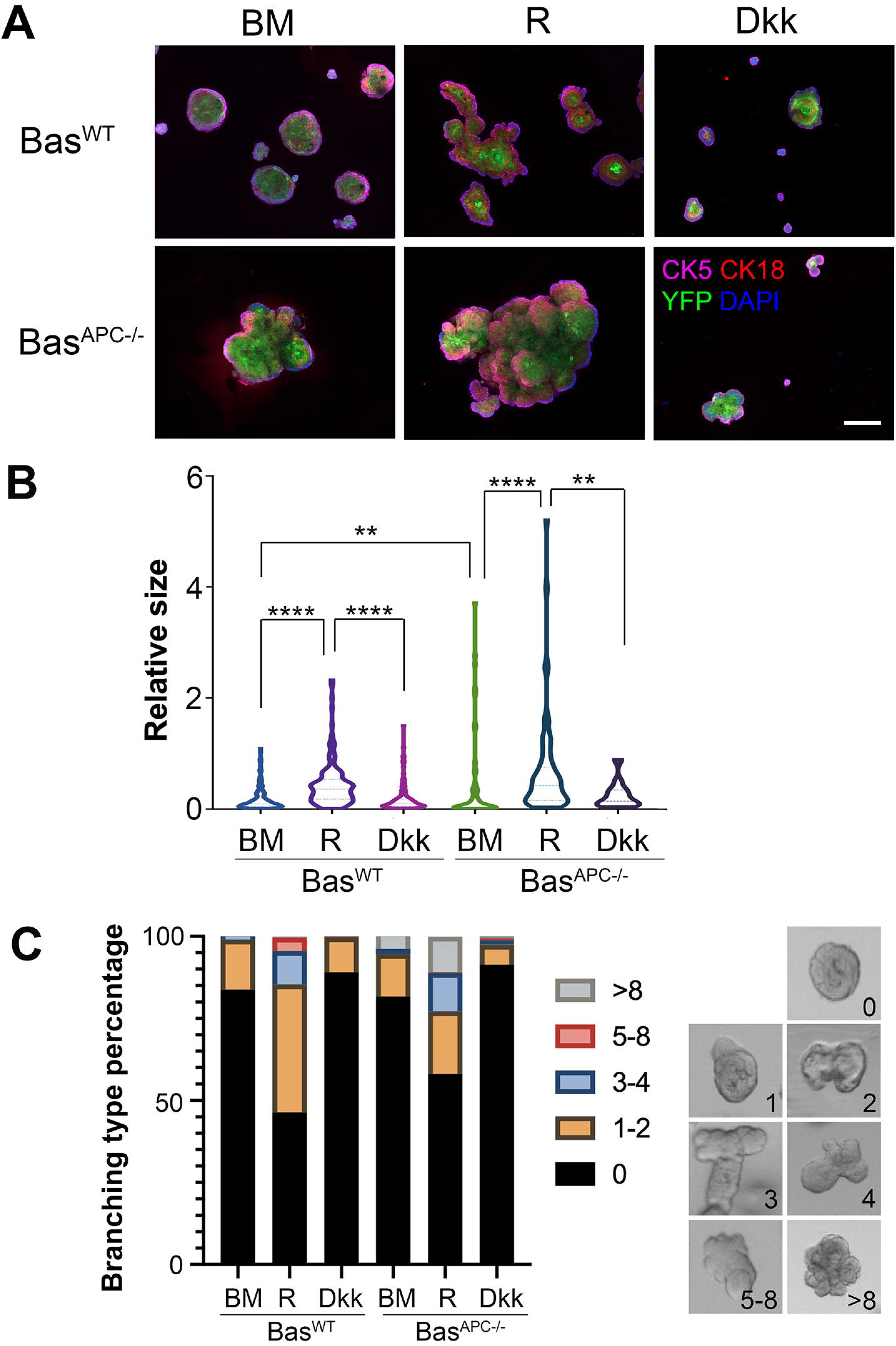
Analysis of organoids derived from basal cells of different Wnt activities. (**A**) Representative IF images showing *Bas*^*WT*^ and *Bas*^*APC−/−*^ organoid morphology after 7 days under different culturing conditions. Scale bar, 200 μm. (**B**) Violin plot comparing relative sizes of *Bas*^*WT*^ and *Bas*^*APC−/−*^ organoids under different culturing conditions by Mann-Whitney U-test. **** *p*<0.0001, ** *p*<0.01. (**C**) Quantification of *Bas*^*WT*^ and *Bas*^*APC−/−*^ organoid branching under different conditions. Examples of different numbers of branching are shown on the right.

### DHT suppresses prostate basal stem cell activities in organoid assay

Given the pivotal role of androgen signaling in prostate development, we next tested how adding DHT affects organoid growth from wild-type basal cells. The basement medium was supplemented with DHT at two concentrations 10^−7^ M and 10^−9^ M as the DHT-9 and DHT-7 conditions. In another two conditions, DHT was also added together with R-spondin (RD-9 and RD-7 conditions) to test the combinatory effects of activating Wnt and androgen signaling pathways on prostate organoid growth. Representative images of organoids growing for 7 days under these conditions are shown in Fig. 3A. Surprisingly, our data consistently showed that DHT suppressed basal stem cell activities in organoid culture in a concentration-dependent manner. When organoid size and structural complexity as measured by the number of branching were quantified, DHT-9 condition led to significantly reduced growth and branching morphogenesis compared to BM alone, and DHT-7 condition further exacerbated such phenotypes as the organoids were significantly stunted in growth (Fig. 3A-C). Adding R-spondin was able to rescue these phenotypes to some extent, but DHT could clearly antagonize the growth-promoting effects of R-spondin since organoids under the RD-9 condition were significantly smaller and contained less branching than those under the R condition (Fig. 3B, 3C). These data imply that Wnt and androgen signaling pathways have opposite effects on prostate basal stem cell activities and that their combined effects may be subtractive.

**Figure 3.**
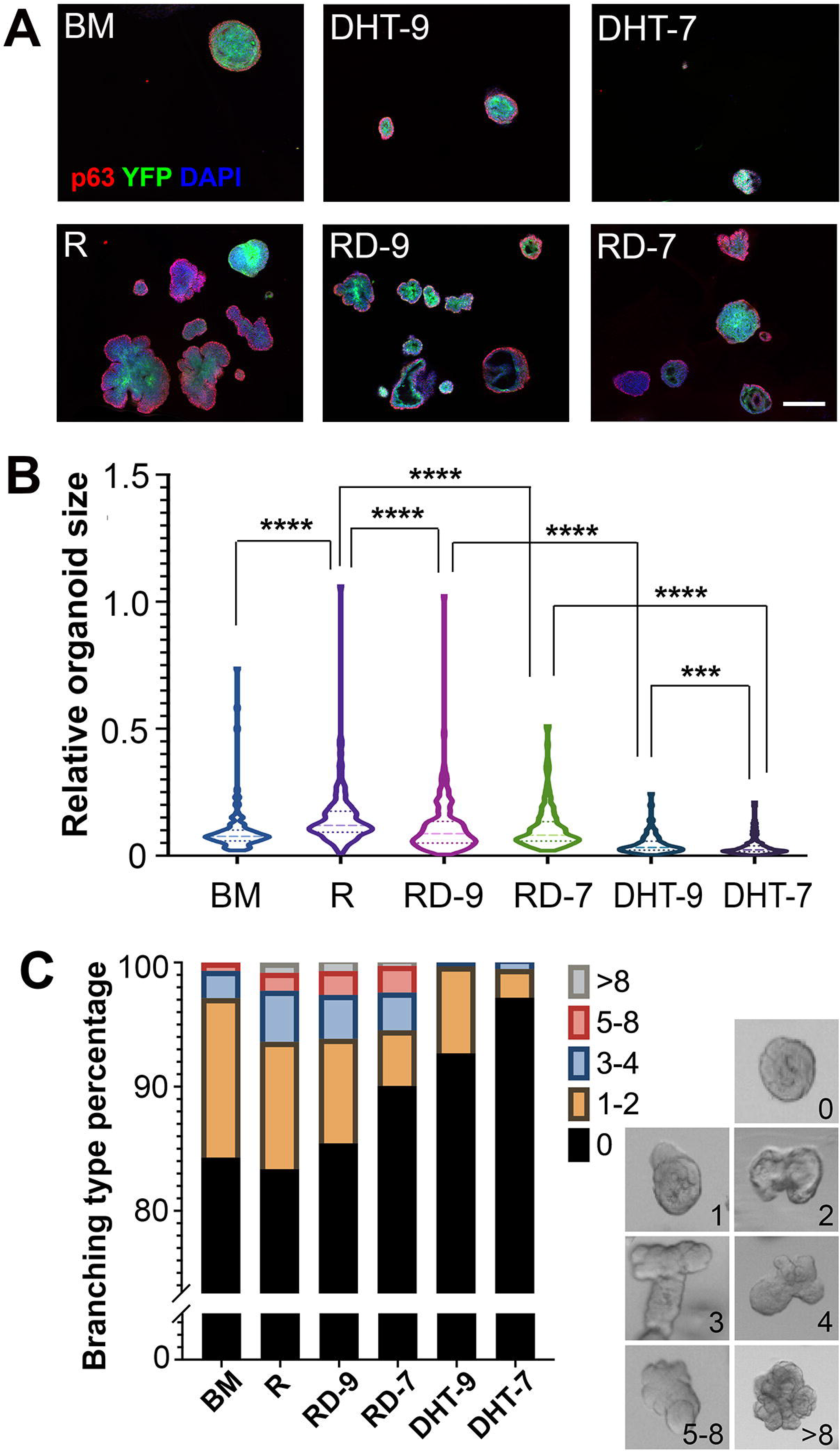
Analysis of basal-derived organoids under combined R-spondin and DHT treatment. (**A**) Representative IF images showing organoid morphology after 7 days under different R-spondin and DHT conditions. Scale bar, 200 μm. (**B**) Violin plot comparing relative organoid sizes under different treatments by Mann-Whitney U-test. **** *p*<0.0001, *** *p*<0.001. (**C**) Quantification of organoid branching under different R-spondin and DHT conditions. Examples of different numbers of branching are shown on the right.

### DHT decreases Wnt signaling activity in prostate organoids

Based on the results above, we hypothesized that DHT inhibits basal organoid growth through modulation of the Wnt signaling pathway. To test it, we isolated basal cells from the *TCF/Lef-H2B.GFP* reporter mice, in which GFP signal can serves as readout for canonical Wnt signaling activity (Ferrer-Vaquer et al., 2010). We then cultured these cells as organoids using 6 medium conditions: BM, R, RD-9, RD-7, DHT-9, and DHT-7. After 7 days of culture, we found that the GFP signals were usually present in the center of the organoids, and were most prominent in the R group (Fig. 4A). When we quantified the percentages of GFP area size of the total organoid size, we found the trend as follows: R>BM>RD-9>RD-7>DHT-9>DHT-7, and adding DHT significantly reduced GFP area even in the presence of R-spondin (Fig. 4B). Expression reduction upon DHT treatment was also observed for several Wnt target genes *Axin2*, *Lef1*, *Ccnd2*, *Cd44*, and *Myc*, as quantified by qRT-PCR (Fig. 4C), confirming that DHT decreases Wnt signaling activity.

**Figure 4.**
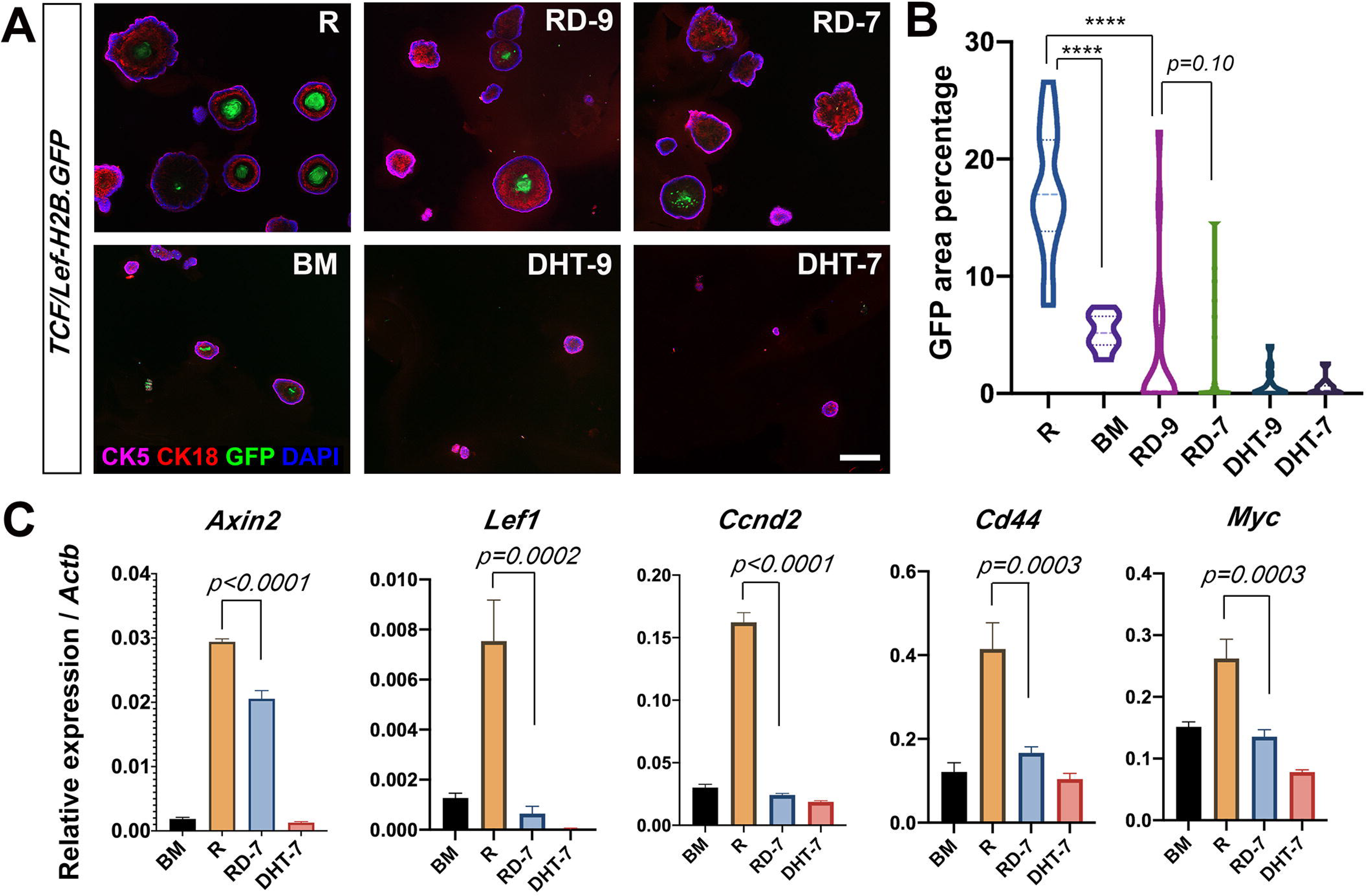
DHT decreases Wnt signaling activities in organoids. (**A**) Representative IF images showing *TCF/Lef-H2B.GFP* reporter signals in basal-derived organoids cultured under different conditions. Scale bar, 200 μm. (**B**) Violin plot showing GFP area percentages for different treatment conditions. **** *P*<0.0001 by using Mann-Whitney U-test. (**C**) Quantitative real-time PCR analysis of Wnt target genes under different treatments. Gene expression levels were normalized to β-actin expression. *P* values were analyzed by unpaired student’s t-test. Error bars correspond to one s.d.

To further explore the mechanism, we isolated mRNAs from BM, R, and RD-7 organoids and performed RNA-sequencing. Principal component analysis (PCA) and unsupervised hierarchical clustering analysis demonstrated the consistency of the samples within each group and the distinct transcriptomes of the RD-7 group from the BM and R groups (Fig. 5A, 5B). Gene expression differential analysis revealed the numbers of upregulated and downregulated genes between each groups as shown in the volcano plots (Fig. S2A). Many genes, including Wnt targets, were upregulated in R compared to BM, but were downregulated in RD-7 compared to R (Fig. S2B). David GO analysis (Huang da et al., 2009) of differentially expressed gene set between R and RD-7 groups identified key pathways involved, among which were cell cycle regulation and Wnt signaling pathway (Fig. 5C), supporting our model that the inhibitory effects of androgen on basal organoids were mediated through modulation of the canonical Wnt activity. In further support of this, gene set enrichment analysis (GSEA) (Subramanian et al., 2005) showed that Wnt signaling pathway genes were highly enriched in the genes that were downregulated in the RD-7 group compared to the R group (Fig. 5D). Similarly, genes of the GO term Organ Morphogenesis were also highly enriched in the downregulated RD-7 genes, while two gene signatures that were previously shown to be upregulated by androgen (Mulholland et al., 2011; Schaeffer et al., 2008) were highly enriched in the upregulated RD-7 genes (Fig. 5D).

**Figure 5.**
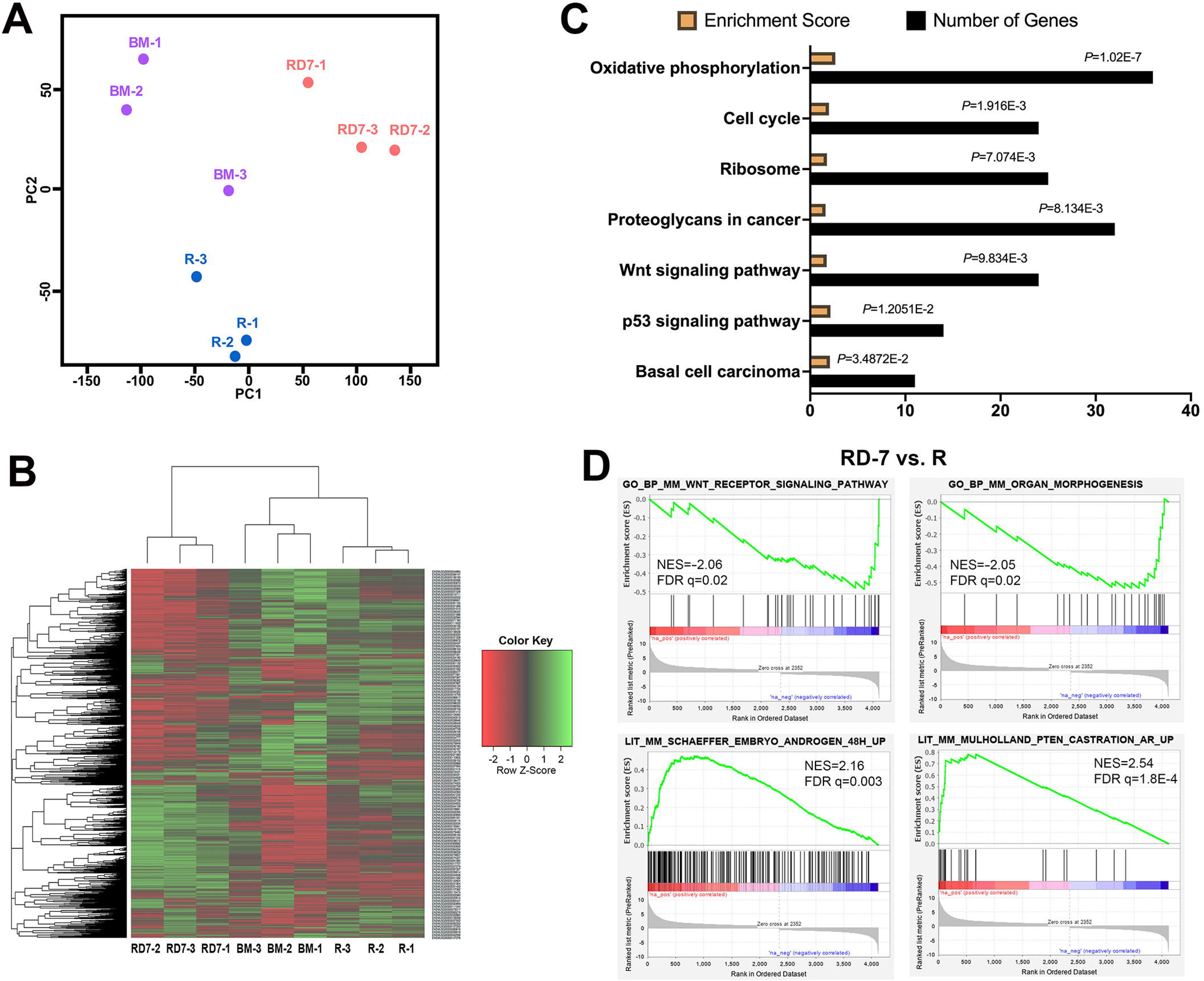
RNA-seq analysis of basal-derived organoids under different treatment conditions. (**A**) Scatter-plot of the two main components from a principle component analysis of the BM (purple), R (blue), and RD-7 (orange) samples. N=3 in each group. (**B**) Unsupervised hierarchical clustering analysis showing distinct transcriptomes of RD-7 from BM and R. (**C**) David GO analysis showing the enriched pathways in the comparison of R vs. RD-7, with the number of genes in each pathway. (**D**) GSEA comparison of RD-7 vs. R showing Wnt pathway genes and organ morphogenesis genes are enriched in the downregulated genes in the RD-7 group, and androgen responsive genes enriched in the upregulated genes in the RD-7 group.

### Increased AR and β-catenin binding in organoid cells upon R-spondin and DHT treatment

Previous studies in prostate cancer cell lines have shown that AR can compete with TCF for β-catenin binding (Song et al., 2003), and the binding of AR and β-catenin can enhance AR signaling output (Amir et al., 2003; Mulholland et al., 2002; Truica et al., 2000). How this binding affects Wnt signaling activity is unclear. We hypothesize that DHT inhibition of Wnt activity in prostate organoids may be mediated through AR binding to β-catenin in the nucleus to hamper the normal function of β-catenin as a Wnt signaling transcription factor. To visualize their interactions in organoids, we performed Duolink Proximity Ligation Assay (PLA) (Fredriksson et al., 2002; Soderberg et al., 2008), in which antibodies recognizing both AR and β-catenin in close proximity can yield an amplified signal dot in situ. We cultured prostate organoids for 7 days and performed quality check experiments to confirm that PLA signals can only be detected when both AR and β-catenin antibodies were applied (Fig. 6A). Interestingly, when the organoids were cultured under BM, R, RD-9, and RD-7 conditions, we observed significantly more AR and β-catenin protein interactions per nucleus or per cell under the RD treatment conditions (Fig. 6B, 6C), and the interactions increased with the concentration of DHT (Fig. 6C).

**Figure 6.**
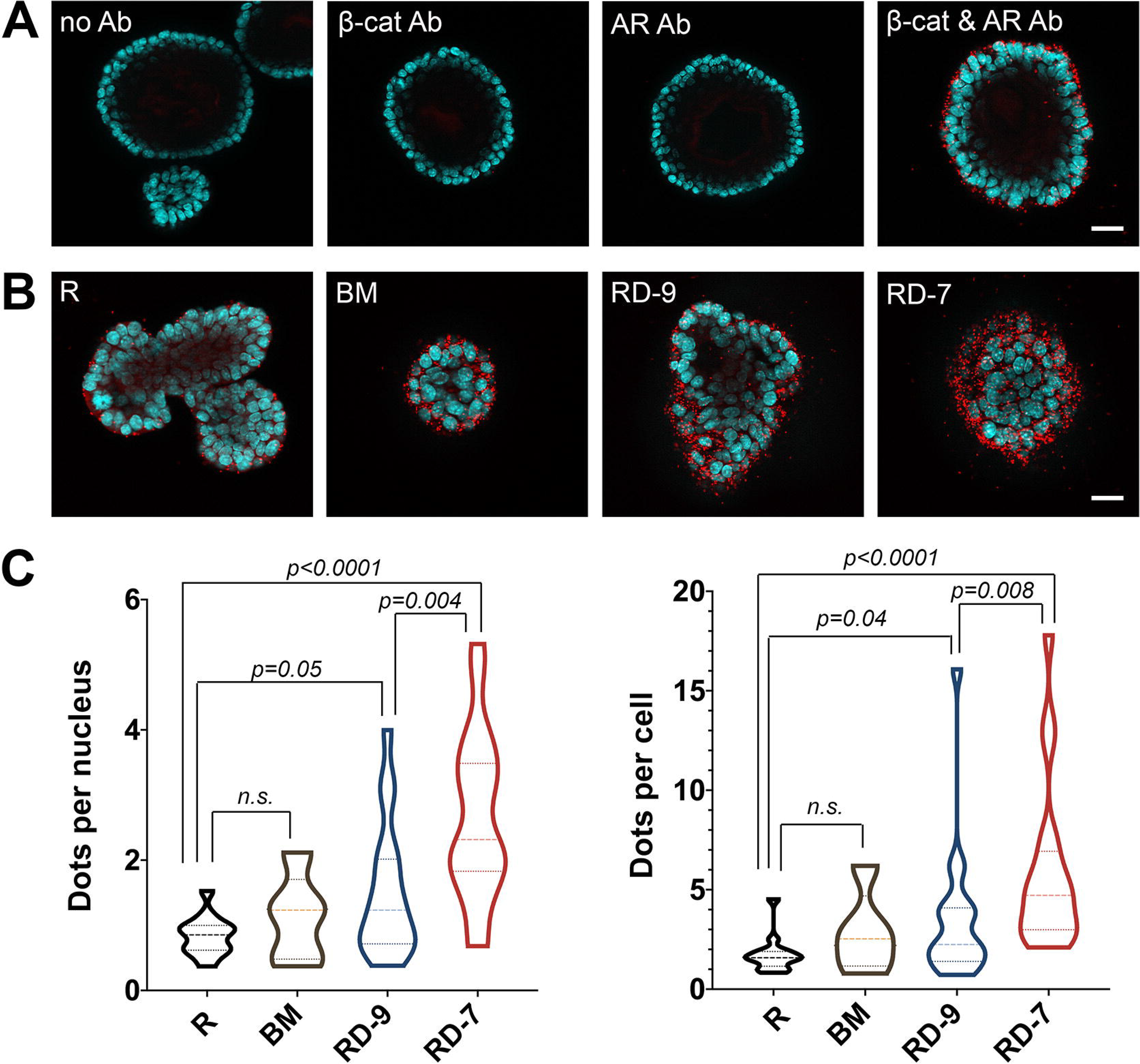
Duolink PLA of AR and β-catenin binding in organoids. (**A**) Representative IF images showing Duolink PLA signals (red dots) only in the presence of both β-catenin and AR antibodies. (**B**) Representative IF images showing Duolink PLA signals under R, BM, RD-9 and RD-7 conditions. Cyan represents nuclear DAPI staining. Scale bars, 50 μm. (**C**) Violin plots showing quantitation of dots per nucleus (left) and dots per cell (right) in each organoid under different treatment conditions. *P* values were analyzed by Mann-Whitney U-test.

These data are consistent with our hypothesis and call for further functional investigation of the consequences of increased AR and β-catenin binding in prostate organoids. Previous studies have only focused on the effects of such interactions on androgen signaling output, and cancer cell lines were often used as materials (Amir et al., 2003; Mulholland et al., 2002; Song et al., 2003; Truica et al., 2000). Analyzing the AR and β-catenin cistromes through ChIP-seq under various treatment conditions for the basal-derived organoids should provide mechanistic insights into how Wnt signaling output is modified by enhanced AR binding in the context of normal prostate stem cell function.

## Methods

### Mouse strains and genotyping

The *CK5-CreER*^*T2*^, *Pten*^*flox*^, and *R26R-CAG-YFP* lines were used previously (Xie et al., 2017). The *Ctnnb1*^*flox*^ (Brault et al., 2001) and *APC*^*flox*^ (Cheung et al., 2010) lines were obtained from JAX. The *TCF/Lef-H2B.GFP* line (Ferrer-Vaquer et al., 2010) was obtained from Dr. Hadjantonakis. Animals were maintained in C57BL/6N or mixed background. Genotyping was performed by PCR using tail genomic DNA, with the following primer sequences: *CreER*^*T2*^ allele: 5’-CAGATGGCGCGGCAACACC-3’ and 5’-GCGCGGTCTGGCAGTAAAAAC-3’; *TCF/Lef-H2B.GFP* allele: 5’-ACAACAAGCGCTCGACCATCAC-3’ and 5’-AGTCGATGCCCTTCAGCTCGAT-3’; *Ctnnb1*^*flox*^ allele: 5’-ACTGCCTTTGTTCTCTTCCCTTCTG-3’ and 5’-CAGCCAAGGAGAGCAGGTGAGG-3’; *APC*^*flox*^ allele: 5’-GAGAAACCCTGTCTCGAAAAAA-3’ and 5’-AGTGCTGTTTCTATGAGTCAAC-3’; *R26R-CAG-YFP* allele: 5’-AAGGGAGCTGCAGTGGAGTA-3’ (wild-type forward), 5’-CCGAAAATCTGTGGGAAGTC-3’ (wild-type reverse), 5’-ACATGGTCCTGCTGGAGTTC-3’ (mutated forward), 5’-GGCATTAAAGCAGCGTATCC-3’ (mutated reverse).

### Mouse procedures

For tamoxifen induction, mice were administered 9 mg per 40 g body weight tamoxifen (Sigma) suspended in corn oil by oral gavage once daily for 4 consecutive days. For BrdU incorporation assay, BrdU (Sigma) was dissolved in PBS (10 mg/ml) and administered by intraperitoneal injection twice daily (0.1 ml per dose) for 11 consecutive days during homeostasis to label proliferating cells. All animal experiments received approval from the Institutional Animal Care and Use Committee at UCSC.

### Tissue collection and flow cytometry

Mouse prostate tissues were dissected and fixed in 4% paraformaldehyde for subsequent cryo-embedding in OCT compound (Sakura), or fixed in 10% formalin followed by paraffin embedding. For flow cytometry, prostate tissues were dissected and minced to small clumps, followed by enzymatic dissociation with 0.2% Collagenase/Hyaluronidase (StemCell Technologies) in DMEM/F12 media with 5% FBS for 3 h at 37°C. Tissues were digested with 0.25% Trypsin-EDTA (StemCell Technologies) for 1 h at 4°C, passed through 21- to 26-gauge syringes and filtered through a 40-mm cell strainer to obtain single-cell suspensions. Dissociated prostate cells were suspended in Hanks’ Balanced Salt Solution Modified/2% FBS. Dead cells were excluded by propidium iodide staining. Lineage-marked basal cells were sorted based on YFP positivity. Antibodies (Table S1) were used for sorting Lin^−^Sca-1^+^ CD49f^hi^ basal cells from *TCF/Lef-H2B.GFP* mice. Cell sorting was performed on a BD FACS Aria II instrument in the Flow Cytometry Shared Facility of UCSC.

### Renal grafting assay

Flow-sorted *APC*^*fl/fl*^ YFP^+^ basal cells were mixed with 2.5 × 10^5^ dissociated urogenital sinus mesenchyme (UGM) cells from embryonic day 18.0 rat embryos. UGM cells were obtained from dissected urogenital sinus treated for 30 min in 1% trypsin, followed by mechanical dissociation and treatment with 0.1% collagenase B (Roche) for 30 min at 37°C, and washing in PBS. Pelleted cell mixtures were resuspended in 10 μl of 1:9 collagen/setting buffer (10× Earle’s Balanced Salt Solution (Life Technologies), 0.2 M NaHCO_3_ and 50 mM NaOH), and gelatinized at 37°C for 20 min. Tissue recombinants were cultured in DMEM/10% FBS supplemented with 10^−7^ M DHT overnight, followed by transplantation under the kidney capsules of immunodeficient NCRNU-M sp/sp nude mice (Taconic Biosciences). Grafts were collected after 3 months of growth and embedded in OCT.

### Prostate organoids culture

Flow-sorted basal cells were washed with advanced DMEM/ F12 (Life Technologies), and resuspended in 10 μl advanced DMEM/F12 and 30 μl Matrigel per well in the Nunc Lab-Tek II CC2 Chamber Slide System (Thermo Fisher Scientific). Chamber slide was put upside down in the 37°C cell culture incubator for 15 min to let the matrigel solidify. Mouse prostate organoid culture medium was prepared according to a previous protocol (Drost et al., 2016). Briefly, the following components were added to advanced DMEM/F12 medium: B27 (50× diluted), HEPES 1 M (100× diluted), GlutaMAX (100× diluted), Penicillin-streptomycin (100× diluted), N-acetylcysteine (1.25 mM), EGF (50 ng/ml), A83-01 (200 nM), Noggin (100 ng/ml), Y-27632 dihydrochloride (10 μM) as basement medium (BM). R-spondin1 (500 ng/ml) and different concentrations of DHT (10^−7^M, 10^−9^M) were added alone or in combinations to BM as different treatments. Dkk-1 (GenScript) was added to BM at final concentration of 100 ng/ml. Organoid culture medium was pre-warmed before adding to the wells. The medium was changed every 3 days. Organoids were fixed in 4% PFA for 20 min at room temperature before immunofluorescence staining. In situ organoid images were taken using the Keyence microscope, and immunofluorescence images were taken using Zeiss AxioImager microscope in the UCSC Microscopy Shared Facility. Organoid sizes and GFP area percentage were quantified using ImageJ.

### Histology and immunofluorescence staining

H&E staining was performed using standard protocols as previously described, and visualized using a Zeiss AxioImager. Immunofluorescence staining was performed using 6 μm cryosections or on organoids in situ. Samples were incubated with 10% normal goat serum (NGS) and primary antibodies diluted in 10% NGS overnight at 4°C. Samples were then incubated with secondary antibodies (diluted 1:500 in PBST) labeled with Alexa Fluor 488, 555, or 647 (Invitrogen/Molecular Probes). Slides were mounted with VectaShield mounting medium with DAPI (Vector Labs), and images were taken on a Leica TCS SP5 spectral confocal microscope in the UCSC Microscopy Shared Facility. All primary antibodies and dilutions used are listed in Table S1.

### Quantitative real-time PCR

Cultured organoids in matrigel were scraped from the incubation chamber with spoon to exclude fibroblasts contamination. Organoids mRNA was isolated using the RNeasy Micro Kit (Qiagen). RNA was reverse transcribed and amplified into cDNA using SuperScrip III kit (Life Technology). Quantitative real-time PCR was carried out using Power SYBR Green PCR Master Mix (Life Technology) in the ViiA 7 Real-Time PCR instrument. Expression values were obtained using the ΔΔCT method and normalized to β-actin (Actb) expression; average values are shown as mean±s.d. Primer sequences are provided in Table S2.

### Duolink proximity ligation assay

Organoids were fixed in 4% PFA for 20 min at room temperature before performing Duolink PLA. Assay was performed as per the kit instruction (DUO92101, Sigma). The antibodies used were AR (Sigma A9853, Rabbit, 1:500 dilution) and β-catenin (BD Bioscience 610153, mouse, 1:500 dilution). Images were taken using the Zeiss AxioImager microscope in the UCSC Microscopy Shared Facility.

### Lineage Analysis and Statistics

For lineage-tracing analysis, cell numbers were counted manually using confocal ×40 photomicrographs across tissue sections. Basal cells were determined based on positive CK5 staining and location at the basement membrane. Luminal cells were determined based on positive CK18 staining, the columnar shape, and location at the apical side of the epithelium. Statistical analyses for lineage tracing and organoids experiments were performed using the two-sided student’s t-test or Mann-Whitney U test as appropriate. At least three biological replicates for each experiment or genotype were analyzed. The variances were similar between the groups that were being statistically compared.

### Organoid bulk RNA-seq

Organoid mRNA was isolated using the RNeasy Micro Kit (Qiagen). RNA was reverse transcribed and amplified into cDNA using the Takara SmartSeq kit at the UC Berkeley QB3 Genomics Center, where library construction and sequencing were performed. The 2×150 bp paired-end sequencing was performed on the NovaSeq 6000 platform. bc and bcl2fastq was used for converting BCL to fastq format, coupled with adaptor trimming. Sequencing reads were then mapped to mouse genome (mm10) using the STAR package. Mapped sequencing reads were assigned to genes using ‘featurecounts’ function of Rsubread package (version 1.30.7). Expression of genes was measured by calculating fragments per kilobase of exon model per million mapped reads (FPKM value) using edgeR package (version 3.24.1) with default settings.

### Principal components analysis and clustering analysis

Genes with extremely low or high expression (the mean FPKM in all samples < 0.3 or > 6000) were filtered out to decrease data noise and potential outliers. The log-transformation was performed on the data. PCA was then performed on the data with ‘prcomp’ function in R v3.5.0 with the parameter scale. = TRUE. The gene hierarchical clustering was done by using ‘heatmap.2’ function of gplots package (version 3.0.1 in R v3.5.0). Here, the Spearman correlation distance was calculated and the complete linkage clustering algorithm was chosen.

### Gene expression and pathway analyses

Differential expression was estimated using the empirical Bayes methods (edgeR v3.24.1 in R v3.5.0) to obtain false discovery rate (FDR) and fold change. The differentially expressed genes (FDR < 0.05, and fold change > 2) were extracted and fed to the DAVID website for the enriched pathway analysis.

### Gene set enrichment analysis

The significantly differentially expressed genes (FDR < 0.05) were ranked by their log-transformed fold change value. Gene Set Enrichment Analysis (GSEA) was conducted using GSEA software (Version 4.0.3). The pre-ranked gene list and MousePath_All_gmt-Format.gmt or MousePath_GO_gmt.gmt gene set (both were downloaded from http://ge-lab.org/gskb/) were used for running the tool “Run GSEA Preranked” with default parameters.

## Acknowledgements

We thank the microscopy and FACS shared facilities at UCSC for technical support. This work was supported by a post-doctoral fellowship from the TRDRP (Y.L.), a Santa Cruz Cancer Benefit Group award (Z.A.W.), and the NIH grant R01GM116872 (Z.A.W.).

## Author contributions

Y.L. and Z.A.W. designed the study. Y.L. performed organoid experiments. C.H. performed lineage tracing and renal grafting experiments. J.W., S.K., and T.C. performed bioinformatic analyses. All the authors discussed data, and contributed to figures and tables. Z.A.W. wrote the manuscript with input from Y.L., J.W., and C.H.

---

**Figure S1. Analysis of *Bas*^*bcat−/−*^ basal cells.**

(**A**) IF images showing β-catenin expression pattern in *Bas*^*bcat−/−*^ mice. Scale bar, 20 μm. (**B**) FACS plot of sorting YFP+ *Bas*^*bcat−/−*^ basal cells. (**C**) qRT-PCR showing significantly reduced β-catenin expression in *Bas*^*bcat−/−*^ basal cells compared to wildtype basal cells by student’s t-test. Gene expression levels were normalized to β-actin expression. (**D**) A mouse losing hair due to β-catenin deletion by the *CK5-CreER*^*T2*^ driver expression in skin basal cells.

**Figure S2. Comparison of organoid RNA-seq data of different treatments.**

**(A)** Volcano plots showing the numbers of upregulated and downregulated genes between R vs. BM, RD-7 vs. R, and BM vs. RD-7 (FDR<0.05 and fold change >2). **(B)** Heatmap showing expression levels of selected genes in different samples.

## References

Amir, A.L., Barua, M., McKnight, N.C., Cheng, S., Yuan, X., and Balk, S.P. (2003). A direct beta-catenin-independent interaction between androgen receptor and T cell factor 4. The Journal of biological chemistry 278, 30828–30834.

Binnerts, M.E., Kim, K.A., Bright, J.M., Patel, S.M., Tran, K., Zhou, M., Leung, J.M., Liu, Y., Lomas, W.E., 3rd, Dixon, M., et al. (2007). R-Spondin1 regulates Wnt signaling by inhibiting internalization of LRP6. Proceedings of the National Academy of Sciences of the United States of America 104, 14700–14705.

Brault, V., Moore, R., Kutsch, S., Ishibashi, M., Rowitch, D.H., McMahon, A.P., Sommer, L., Boussadia, O., and Kemler, R. (2001). Inactivation of the beta-catenin gene by Wnt1-Cre-mediated deletion results in dramatic brain malformation and failure of craniofacial development. Development 128, 1253–1264.

Carmon, K.S., Gong, X., Lin, Q., Thomas, A., and Liu, Q. (2011). R-spondins function as ligands of the orphan receptors LGR4 and LGR5 to regulate Wnt/beta-catenin signaling. Proceedings of the National Academy of Sciences of the United States of America 108, 11452–11457.

Cheung, A.F., Carter, A.M., Kostova, K.K., Woodruff, J.F., Crowley, D., Bronson, R.T., Haigis, K.M., and Jacks, T. (2010). Complete deletion of Apc results in severe polyposis in mice. Oncogene 29, 1857–1864.

Choi, N., Zhang, B., Zhang, L., Ittmann, M., and Xin, L. (2012). Adult murine prostate basal and luminal cells are self-sustained lineages that can both serve as targets for prostate cancer initiation. Cancer cell 21, 253–265.

Chua, C.W., Shibata, M., Lei, M., Toivanen, R., Barlow, L.J., Bergren, S.K., Badani, K.K., McKiernan, J.M., Benson, M.C., Hibshoosh, H., et al. (2014). Single luminal epithelial progenitors can generate prostate organoids in culture. Nature cell biology.

Clevers, H., Loh, K.M., and Nusse, R. (2014). Stem cell signaling. An integral program for tissue renewal and regeneration: Wnt signaling and stem cell control. Science 346, 1248012.

Cunha, G.R., Alarid, E.T., Turner, T., Donjacour, A.A., Boutin, E.L., and Foster, B.A. (1992). Normal and abnormal development of the male urogenital tract. Role of androgens, mesenchymal-epithelial interactions, and growth factors. Journal of andrology 13, 465–475.

de Lau, W., Barker, N., Low, T.Y., Koo, B.K., Li, V.S., Teunissen, H., Kujala, P., Haegebarth, A., Peters, P.J., van de Wetering, M., et al. (2011). Lgr5 homologues associate with Wnt receptors and mediate R-spondin signalling. Nature 476, 293–297.

Drost, J., Karthaus, W.R., Gao, D., Driehuis, E., Sawyers, C.L., Chen, Y., and Clevers, H. (2016). Organoid culture systems for prostate epithelial and cancer tissue. Nature protocols 11, 347–358.

Ferrer-Vaquer, A., Piliszek, A., Tian, G., Aho, R.J., Dufort, D., and Hadjantonakis, A.K. (2010). A sensitive and bright single-cell resolution live imaging reporter of Wnt/ss-catenin signaling in the mouse. BMC developmental biology 10, 121.

Francis, J.C., Thomsen, M.K., Taketo, M.M., and Swain, A. (2013). beta-catenin is required for prostate development and cooperates with Pten loss to drive invasive carcinoma. PLoS genetics 9, e1003180.

Fredriksson, S., Gullberg, M., Jarvius, J., Olsson, C., Pietras, K., Gustafsdottir, S.M., Ostman, A., and Landegren, U. (2002). Protein detection using proximity-dependent DNA ligation assays. Nature biotechnology 20, 473–477.

Glinka, A., Wu, W., Delius, H., Monaghan, A.P., Blumenstock, C., and Niehrs, C. (1998). Dickkopf-1 is a member of a new family of secreted proteins and functions in head induction. Nature 391, 357–362.

Holland, J.D., Klaus, A., Garratt, A.N., and Birchmeier, W. (2013). Wnt signaling in stem and cancer stem cells. Current opinion in cell biology 25, 254–264.

Huang da, W., Sherman, B.T., and Lempicki, R.A. (2009). Bioinformatics enrichment tools: paths toward the comprehensive functional analysis of large gene lists. Nucleic acids research 37, 1–13.

Julio, M.K., Shibata, M., Desai, N., Reynon, M., Halili, M.V., Hu, Y.P., Price, S.M., Abate-Shen, C., and Shen, M.M. (2013). Canonical Wnt signaling regulates Nkx3.1 expression and luminal epithelial differentiation during prostate organogenesis. Developmental dynamics: an official publication of the American Association of Anatomists.

Karthaus, W.R., Iaquinta, P.J., Drost, J., Gracanin, A., van Boxtel, R., Wongvipat, J., Dowling, C.M., Gao, D., Begthel, H., Sachs, N., et al. (2014). Identification of Multipotent Luminal Progenitor Cells in Human Prostate Organoid Cultures. Cell.

Kishimoto, J., Burgeson, R.E., and Morgan, B.A. (2000). Wnt signaling maintains the hair-inducing activity of the dermal papilla. Genes & development 14, 1181–1185.

Lai, K.P., Yamashita, S., Vitkus, S., Shyr, C.R., Yeh, S., and Chang, C. (2012). Suppressed prostate epithelial development with impaired branching morphogenesis in mice lacking stromal fibromuscular androgen receptor. Mol Endocrinol 26, 52–66.

Lu, T.L., and Chen, C.M. (2015). Differential requirements for beta-catenin in murine prostate cancer originating from basal versus luminal cells. The Journal of pathology 236, 290–301.

Mulholland, D.J., Cheng, H., Reid, K., Rennie, P.S., and Nelson, C.C. (2002). The androgen receptor can promote beta-catenin nuclear translocation independently of adenomatous polyposis coli. The Journal of biological chemistry 277, 17933–17943.

Mulholland, D.J., Tran, L.M., Li, Y., Cai, H., Morim, A., Wang, S., Plaisier, S., Garraway, I.P., Huang, J., Graeber, T.G., et al. (2011). Cell autonomous role of PTEN in regulating castration-resistant prostate cancer growth. Cancer cell 19, 792–804.

Ousset, M., Van Keymeulen, A., Bouvencourt, G., Sharma, N., Achouri, Y., Simons, B.D., and Blanpain, C. (2012). Multipotent and unipotent progenitors contribute to prostate postnatal development. Nature cell biology 14, 1131–1138.

Schaeffer, E.M., Marchionni, L., Huang, Z., Simons, B., Blackman, A., Yu, W., Parmigiani, G., and Berman, D.M. (2008). Androgen-induced programs for prostate epithelial growth and invasion arise in embryogenesis and are reactivated in cancer. Oncogene 27, 7180–7191.

Simons, B.W., Hurley, P.J., Huang, Z., Ross, A.E., Miller, R., Marchionni, L., Berman, D.M., and Schaeffer, E.M. (2012). Wnt signaling though beta-catenin is required for prostate lineage specification. Developmental biology 371, 246–255.

Soderberg, O., Leuchowius, K.J., Gullberg, M., Jarvius, M., Weibrecht, I., Larsson, L.G., and Landegren, U. (2008). Characterizing proteins and their interactions in cells and tissues using the in situ proximity ligation assay. Methods 45, 227–232.

Song, L.N., Herrell, R., Byers, S., Shah, S., Wilson, E.M., and Gelmann, E.P. (2003). Beta-catenin binds to the activation function 2 region of the androgen receptor and modulates the effects of the N-terminal domain and TIF2 on ligand-dependent transcription. Molecular and cellular biology 23, 1674–1687.

Subramanian, A., Tamayo, P., Mootha, V.K., Mukherjee, S., Ebert, B.L., Gillette, M.A., Paulovich, A., Pomeroy, S.L., Golub, T.R., Lander, E.S., et al. (2005). Gene set enrichment analysis: a knowledge-based approach for interpreting genome-wide expression profiles. Proceedings of the National Academy of Sciences of the United States of America 102, 15545–15550.

Toivanen, R., and Shen, M.M. (2017). Prostate organogenesis: tissue induction, hormonal regulation and cell type specification. Development 144, 1382–1398.

Truica, C.I., Byers, S., and Gelmann, E.P. (2000). Beta-catenin affects androgen receptor transcriptional activity and ligand specificity. Cancer research 60, 4709–4713.

Wang, Z.A., Mitrofanova, A., Bergren, S.K., Abate-Shen, C., Cardiff, R.D., Califano, A., and Shen, M.M. (2013). Lineage analysis of basal epithelial cells reveals their unexpected plasticity and supports a cell-of-origin model for prostate cancer heterogeneity. Nature cell biology 15, 274–283.

Wang, Z.A., Toivanen, R., Bergren, S.K., Chambon, P., and Shen, M.M. (2014). Luminal Cells Are Favored as the Cell of Origin for Prostate Cancer. Cell reports.

Xie, Q., Liu, Y., Cai, T., Horton, C., Stefanson, J., and Wang, Z.A. (2017). Dissecting cell-type-specific roles of androgen receptor in prostate homeostasis and regeneration through lineage tracing. Nature communications 8, 14284.

Xin, L., Ide, H., Kim, Y., Dubey, P., and Witte, O.N. (2003). In vivo regeneration of murine prostate from dissociated cell populations of postnatal epithelia and urogenital sinus mesenchyme. Proceedings of the National Academy of Sciences of the United States of America 100 Suppl 1, 11896–11903.

