## Supplementary figures and images for "Modulation of the canonical Wnt activity by androgen signaling in prostate epithelial basal stem cells"

### Supplemental Figure S1

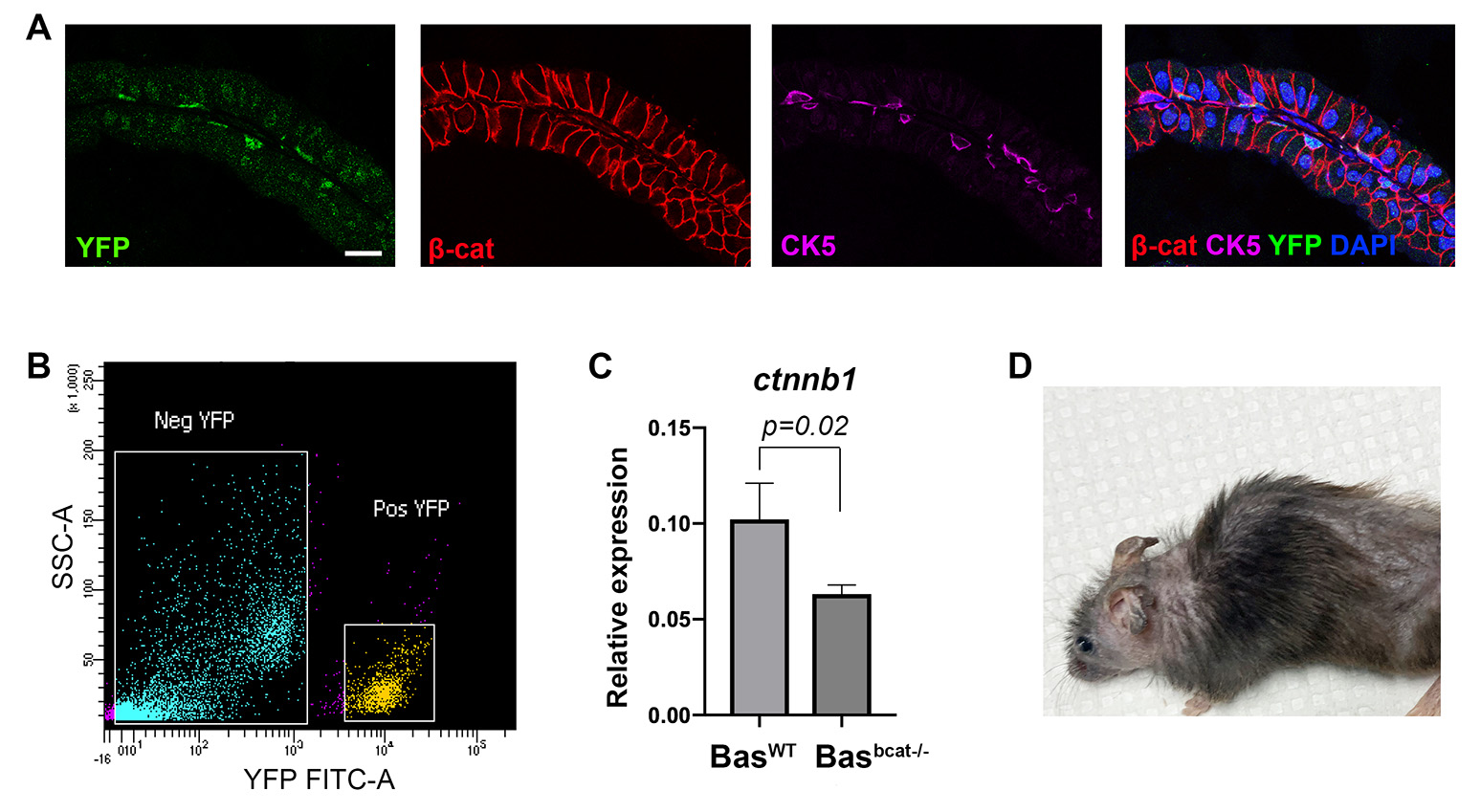

### Supplemental Figure S2

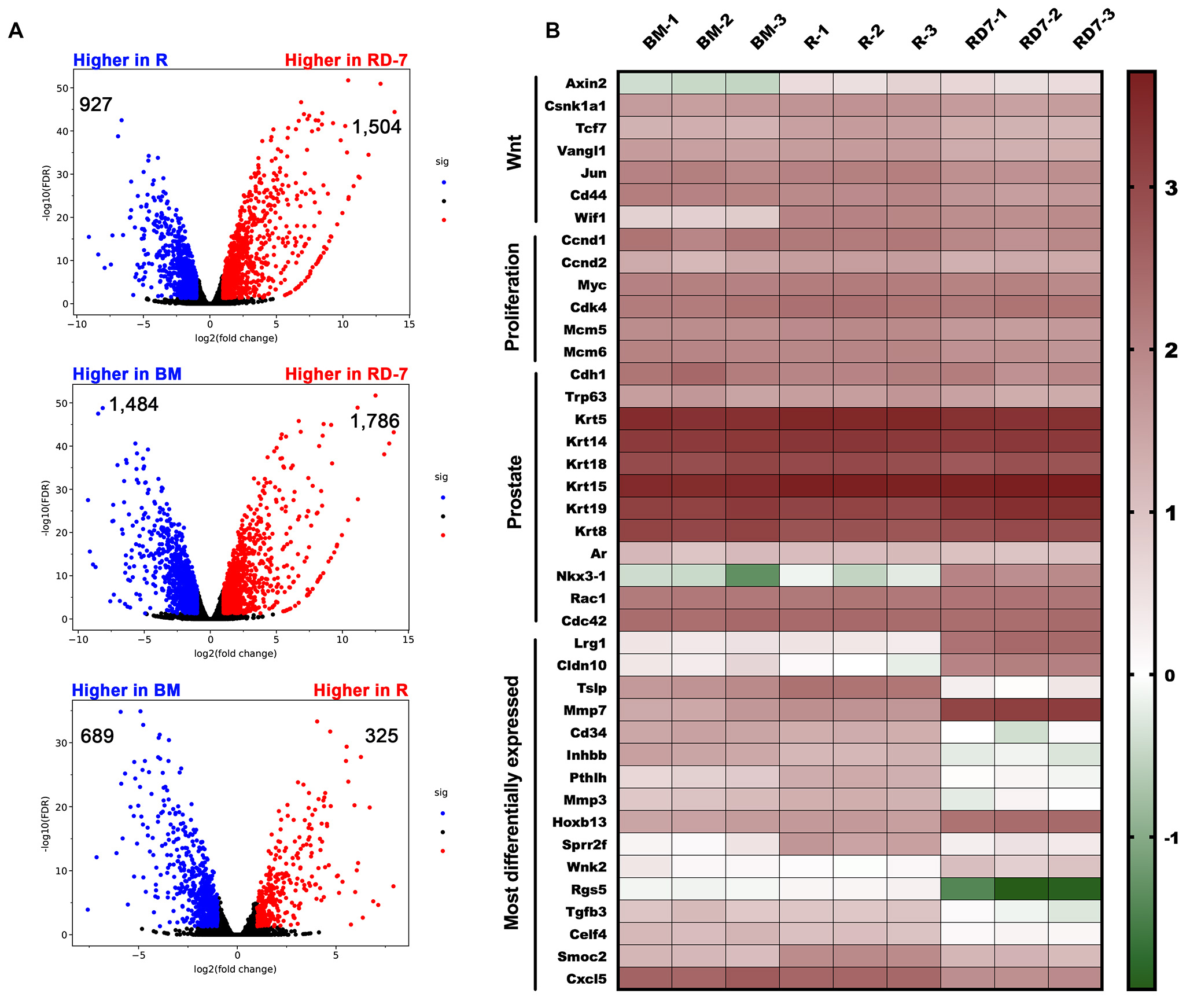
