## Supplemental Table S1 for "Modulation of the canonical Wnt activity by androgen signaling in prostate epithelial basal stem cells"

**Table S1. Antibodies used in this study**

Antibodies for flow cytometry

| Antibody | Supplier | Dilution |
|---|---|---|
| Sca-1-PE-Cy7 | Biolegend clone E13-<br>161.7 #122513 | 1:500 |
| CD49f-PE | eBiosciences clone<br>eBioGoH3 #12-0495 | 1:300 |
| Ter119-eFluor450 | eBiosciences clone Ter119 #48-5921 | 1:250 |
| CD31-eFluor450 | eBiosciences clone 390<br>#48-0311 | 1:250 |
| CD45-eFluor450 | CD45-eFluor450eBiosciences clone<br>30-<br>F11 #48-0451 | 1:250 |

Primary antibodies used for immunofluorescence staining and duolink PLA

| Antigen | Supplier | Ig type | Dilution |
|---|---|---|---|
| BrdU | Serotec #MCA2060 | Rat IgG2a | 1:500 |
| Ki67 | DakoCytomation<br>#M7249 | Rat IgG2a | 1:600 |
| YFP | Abcam #13970 | Chick IgY | 1:2000 |
| CK5 | Covance #PRB-160P | Rabbit IgG | 1:1000 |
| CK18 | Abcam #ab668 | Mouse IgG1 | 1:200 |
| p63 | GeneTex #GTX102425 | Rabbit IgG | 1:1000 |
| AR | Sigma #A9853 | Rabbit IgG | 1:500 |
| β-catenin | BD Biosciences<br>#610153 | Mouse IgG1 | 1:500 for<br>Duolink<br>1:1000 for IF |
