## Supplemental Table S2 for "Modulation of the canonical Wnt activity by androgen signaling in prostate epithelial basal stem cells"

**Table S2. Primers for qPT-PCR**

| Genes | Primer sequence | |
|---|---|---|
| Axin2 | forward | 5’-CAGAGGGACAGGAACCACTC-3’ |
| | reverse | 5’-TGGACACTTGCCAGTTTCTT-3’ |
| Lef1 | forward | 5’-ACGACAAGGCCAGAGAACA-3’ |
| | reverse | 5’-GTCGCTGTTCATATTGGGCA-3’ |
| Ccnd2 | forward | 5’-AAGGACCGGTGCGAGTCA-3’ |
| | reverse | 5’-GGGAGTGCTTCCCTTACCTC-3’ |
| Cd44 | forward | 5’-TCGATTTGAATGTAACCTGCCG-3’ |
| | reverse | 5’-CAGTCCGGGAGATACTGTAGC-3’ |
| Myc | forward | 5’-CCCTAGTGCTGCATGAGGA-3’ |
| | reverse | 5’-TGCCTCTTCTCCACAGACAC-3’ |
| Ctnnb1 | forward | 5’-CATCCCACTGGCCTCTGATA-3’ |
| | reverse | 5’-TCGTGGAATAGCACCCTGTT-3’ |
| Actb | forward | 5’-CGCCACCAGTTCGCCATGGA-3’ |
| | reverse | 5’-TACAGCCCGGGGAGCATCGT-3’ |
